# A watershed model of individual differences in fluid intelligence

**DOI:** 10.1101/041368

**Authors:** Rogier A. Kievit, Simon W. Davis, John Griffiths, Marta Correia, Cam-CAN, Richard N. Henson

## Abstract

Fluid intelligence is a crucial cognitive ability that predicts key life outcomes across the lifespan. Strong empirical links exist between fluid intelligence and processing speed on the one hand, and white matter integrity and processing speed on the other. We propose a watershed model that integrates these three explanatory levels in a principled manner in a single statistical model, with processing speed and white matter figuring as intermediate endophenotypes. We fit this model in a large (N=555) adult lifespan cohort from the Cambridge Centre for Ageing and Neuroscience (Cam-CAN) using multiple measures of processing speed, white matter health and fluid intelligence. The model fit the data well, outperforming competing models and providing evidence for a many-to-one mapping between white matter integrity, processing speed and fluid intelligence. The model can be naturally extended to integrate other cognitive domains, endophenotypes and genotypes.

## 1.1 Introduction

Fluid intelligence, or fluid reasoning, is a core feature of human cognition. It refers to the ability to solve novel, abstract problems that do not depend on task-specific knowledge (Blair, 2006; Carroll, 1993; Deary, 2012; Horn and Cattell, 1966). In contrast to crystallized intelligence, which continues to improve across most of the lifespan, fluid intelligence shows strong age-related declines (Horn and Cattell, 1966; Salthouse, 2009). Understanding the causes of this decline is important for healthy ageing, as preserved fluid intelligence is strongly associated with independent day-to-day functioning (Tucker-Drob, 2011; Willis and Schaie, 1986), and is inversely related to mortality risk (Aichele et al., 2015). At the other end of the lifespan, low fluid intelligence in adolescence predicts poor outcome in later life (Huepe et al., 2011) and is a risk factor for psychopathologies such as schizophrenia (Blair, 2006; Snitz et al., 2006). However, our understanding of how this crucial cognitive ability relates to broader, mechanistic frameworks of cognition and the brain is limited. A promising line of research focuses on the relationships between fluid intelligence, processing speed and white matter organisation. Although intriguing, these empirical relationships are often interpreted in isolation, relating fluid reasoning to processing speed (e.g. Sheppard & Vernon, 2008), processing speed to white matter (e.g. Penke et al., 2010), or fluid intelligence to white matter (e.g. Haász et al., 2013), but never the three together. One unresolved question is therefore whether fluid intelligence, processing speed and white matter can be thought of as part of a single, hierarchical system.

Here, we propose a statistical framework to examine this question, developed by formalizing a conceptual model taken from the literature on psychopathological constructs and their causes. This so-called ‘watershed model’ (Cannon and Keller, 2006) uses the metaphor of a river system to illustrate how complex behavioural traits can be seen as the downstream consequence of many small upstream (e.g., neural/genetic) contributions. From this perspective, the relationship between fluid intelligence (hereafter FI), processing speed (PS) and white matter (WM) is hierarchical, such that WM influences PS, which in turn affects performance on tests of FI. We show that this model naturally accommodates a wide and disparate range of empirical findings, integrates a series of relatively well-established findings into a single larger model, and, most importantly, can be formally tested using Structural Equation Modelling (SEM). We derive a variety of statistical predictions that follow from the watershed model, and use SEM to test these predictions empirically in a large (N=555), population-based sample of ageing adults (18–87 years, Cam-CAN). First, we examine the empirical evidence concerning FI, PS and WM.

### 1.2 Processing speed, fluid intelligence and white matter

Processing speed refers to the general speed with which mental computations are performed. It has been considered a central feature of higher cognitive functioning since the development of the first formalized models of (fluid) intelligence (Salthouse, 1982; Spearman, 1927). It shows comparatively steep age-related declines, similar to or even stronger than FI (Horn & Cattell, 1966; Salthouse, 2000; Schaie, 1994), including in longitudinal samples (Deary and Der, 2005). Processing speed is a broad concept that has can be measured in a variety of ways (Salthouse, 2000). One common approach is to use a set of tasks with strict time-limits, and consider the shared variance across those tasks to reflect an individuals' ability to perform cognitive tasks under time pressure (Babcock et al., 1997). Even purely physiological measures have been considered, such as the latency of neural evoked responses (Salthouse, 2000, Schubert et al., 2015). Other possibilities include the parameters estimated from response time distributions in a single task, such as the mean, standard deviation and exponential for an ex-gaussian distribution, or parameters such as drift rate and boundary separation in diffusion models (Matzke & Wagenmakers, 2009, Ratcliff, Smith, Brown, & McKoon, 2016). Here, we focus on the most basic and simple notions of processing speed, sometimes called psychomotor speed, namely the mean and standard deviation of RT distributions for simple tasks.

The empirical association between PS and FI is one of the most robust findings in psychology (Sheppard and Vernon, 2008). This association holds across the lifespan (Salthouse, 1994), in both healthy elderly (Ritchie et al., 2014) and in the extremes of mental retardation (e.g. Kail, 1992). Longitudinal studies of either end of the lifespan show similar patterns. Dougherty and Haith (1997) showed that infant reaction time at 3.5 months predicts IQ several years later, and Fry and Hale (1996) showed in 214 children and adolescents how longitudinal changes in processing speed mediated changes in fluid intelligence and working memory. At the other end of the lifespan, declines in PS and FI show considerable correlations in old age, with estimates ranging from .53 (Zimprich & Martin, 2002) to .78 (Ritchie et al., 2014). Similarly, a large longitudinal cohort study (Ghisletta, Rabbitt, Lunn, & Lindenberger, 2012) showed that a considerable portion of within-subject age-related decline was shared between FI and PS. Although few studies have explicitly examined the temporal ordering of developmental changes, those that do generally find that declines in PS affect declines in FI and related cognitive abilities. For instance, Kail (2007) examined 185 children (age 8-13) tested twice on multiple outcomes, and found that the best mediation model described a developmental cascade, wherein improvements in processing speed affected working memory which in turn enhanced reasoning. In older adults, Robitaille et al. (2013) showed in two separate cohorts that within-subject declines in processing speed mediated within-subject declines in multiple cognitive domains, including fluid reasoning. Finally, Finkel, Reynolds, McArdle and Pedersen (2007) used bivariate latent change score models in older adults to show that processing speed was a leading indicator of cognitive changes, including in abstract reasoning tasks. Together, these behavioural findings suggest a strong relationship between processing speed and fluid reasoning ability.

The most common metric of PS is the central tendency, such as the mean or median, of RTs on a simple reaction time task. However, individual differences in the *variability* of RTs also relate to fluid reasoning ability (Rabbitt, 1993), such that less variable responses are associated with higher scores on fluid reasoning tasks. This ‘cognitive consistency’ in RTs has been shown to predict cognitive performance in elderly subjects beyond mean RT (MacDonald, Li, &Bäckman, 2009). Both the central tendency and variability of PS predict all-cause mortality (Batterham et al., 2014; Hagger-Johnson et al., 2014), supporting the idea that both are important and independent components of PS. The role of variability can be observed even on the purely neural level: A study using EEG in young adults(Euler et al., 2015) found evidence for the role of variability of neural responses, such that individuals with more stable (less variable) responses to novel stimuli tended to have higher fluid reasoning ability.

Recent work suggests that the proper conceptualisation of the relation between PS and FI is as a causal factor (e.g., Kail, 2000; Rindermann and Neubauer, 2004; Robitaille et al., 2013). The most influential causal account comes from Salthouse (1996), who suggested at least two mechanisms by which PS affects cognitive performance, namely the *limited time mechanism* and the *simultaneity mechanism.* The former suggests that in any timed task, slower speed of processing simply precludes the timely completion of cognitive operations, leading to poorer scores; the latter suggests that high PS is necessary to juggle mental representations simultaneously, in order to perform complex cognitive operations (see Burzynska et al., 2013, for neuroimaging evidence for this claim). More recent work (Schubert et al., 2015) used drift-diffusion and EEG modelling to show that there are multiple components to processing speed, and that these components play different causal roles in different cognitive tasks. In summary, nearly all of the papers reviewed above, either explicitly or implicitly, consider PS to be a ‘lower’, or more fundamental, mental process that is not identical to FI itself (see also Schubert et al., 2015). We can also go further down this presumed causal hierarchy to understand the possible determinants of PS. One such candidate is the structural organization of white matter tracts.

Among the most influential studies showing the importance of white matter organisation are two papers by Penke and colleagues, who showed that the first principal component of fractional anisotropy (FA, a measure of white matter organization) predicted both information processing speed (Penke et al., 2010) as well as general intelligence (Penke et al., 2012). Further work has shown that decreased WM organisation has been associated with decreased PS both in healthy adults (Tuch et al., 2005; Penke et al., 2010) and in individuals suffering from clinical conditions associated with WM loss such as Multiple Sclerosis (Kail, 1997, 1998; Roosendaal et al., 2009; Segura et al., 2010; see Bennett & Madden, 2013, for a review). However, in a sample of 90 older adults, Yang, Bender, & Raz (2014) did not find strong associations between white matter organisation and reaction time components derived from a diffusion model. WM organisation has also been associated with the variability of RTs in children (Tamnes et al., 2012), in healthy controls and preclinical Alzheimer's dementia (Jackson et al. 2012), and decline in WM has been proposed as a key cause of age-related changes in cognition (O'Sullivan et al., 2001). This relationship between WM and performance variability has been found to strengthen with age (Fjell et al., 2011; Laukka et al., 2013; Lövdén et al., 2013b). Other studies have found direct relationships between WM measures and FI (Haász et al., 2013; Kievit et al., 2014) and specific neural (including white matter) structural correlates of intraindividual variability (MacDonald et al., 2009, 2006). Similarly, lesions in WM predict age-related declines in mental speed (Rabbitt et al., 2007a). Assessing a broad set of cognitive and neural markers in a large, age-heterogeneous cohort, Hedden et al. (2014, p. 1) conclude that ‘The largest relationships linked FA and striatum volume to processing speed and executive function’.

A critical link in our model is the behavioural consequence of the microstructural properties evident in the white matter structures, as they are presumed critical for signal transmission between disparate regions of cortex. Various mechanisms (although none demonstrated definitively) have been proposed to explain the relation between WM and PS. One hypothesis is that inefficient signal transmission weakens the signal in neural activity, and/or increases the background noise, ultimately leading to slower decisions (Kail, 1997; Rolls and Deco, 2015). While ageing is generally thought to be accompanied by reduced neuronal plasticity, a growing number of models have addressed the complex—and not uniformly depressing—possibility that age-related changes in brain-behaviour relationships are driven by shifting adaptations to this changing signal-to-noise ratio (Welford, 1984). Such approaches take as a premise the idea that aging reflects a progressive refinement and optimization of generative models used by the brain to predict states of the world (Moran et al.,2014). In this context, observed reductions in axonal density in aging (Peters, 2009) may reflect longterm pruning and adaptation. A related hypothesis that is gaining support is the proposal that age-related demyelination affects propagation of action potentials (Bartzokis et al., 2010), an explanation consistent with slowing in patients with MS (Turken et al., 2008). Unlike MS, however, observations of an age-related increase in dystrophic myelin are relatively rare in macaque microscopy studies, leading Peters (2009) to propose remyelination with shorter internodes as the cause of age-related slowing observed in neurophysiological data. Whilst much further research is needed, these mechanistic accounts help to explain the strong and consistent neurocognitive relationship between PS and WM (Bennett and Madden, 2014; Penke et al., 2010; Turken et al., 2008). Together this suggests a hierarchical relationship, where WM affects PS, which in turn affects FI. Below, we describe a model that can integrate these diverse findings.

## 1.3 Watershed Model

Our goal is to integrate the three explanatory levels (FI, PS and WM) into a single model that can address the range of empirical findings described above. This a general challenge in cognitive neuroscience, namely that of *reductionism* (Kievit et al., 2011a): How do we best relate the phenotype observed at the ‘higher’ level of measurement (e.g., scores on a test of fluid intelligence) to ‘lower’ levels of explanation (e.g., WM structure)? We next show how a theoretical model from the field of psychopathology can be translated into a testable psychometric model to achieve this goal.

In psychopathology, single cause models for mental disorders such as schizophrenia were initially popular, but have not been successful: Despite being highly heritable and having various structural brain correlates, the search for single (or even a limited set of) genetic loci has not yielded candidates that explain more than a trivial percentage of the variance of the phenotypes of interest. Recently, the ‘watershed model’ proposed by Cannon and Keller (2006) has provided a conceptual framework to help understand the potential multiple determinacy between various explanatory levels in the study of mental disorders (see also Penke et al., 2007)^1^.

A simplified representation of the watershed model is shown in *Figure 1.* The central idea is that an observable phenotype can be thought of as the mouth of a river (denoted by “1” in the Figure), and is the end product of a wide range of small, causal, genetic influences (genotypes) that exert their influence through a series of intermediate endophenotypes (such as neural and cognitive variables). A crucial assumption in this model is that genetic influences do not directly affect the phenotype, but do so indirectly via endophenotypes. These endophenotypes are the hypothesized intermediate mechanistic steps between many small genetic influences that together exert considerable influence (the idea of many small genetic effects has been referred to as the Fourth Law of behavioural genetics, c.f. C. F. Chabris, Lee, Cesarini, Benjamin, & Laibson, 2015). Such a model allows us to integrate the disparate known endophenotypes as potentially independent upstream ‘tributaries’ (denoted by “2a”-“2d”) that all contribute to the distal consequence (“1”).

**Figure 1:**
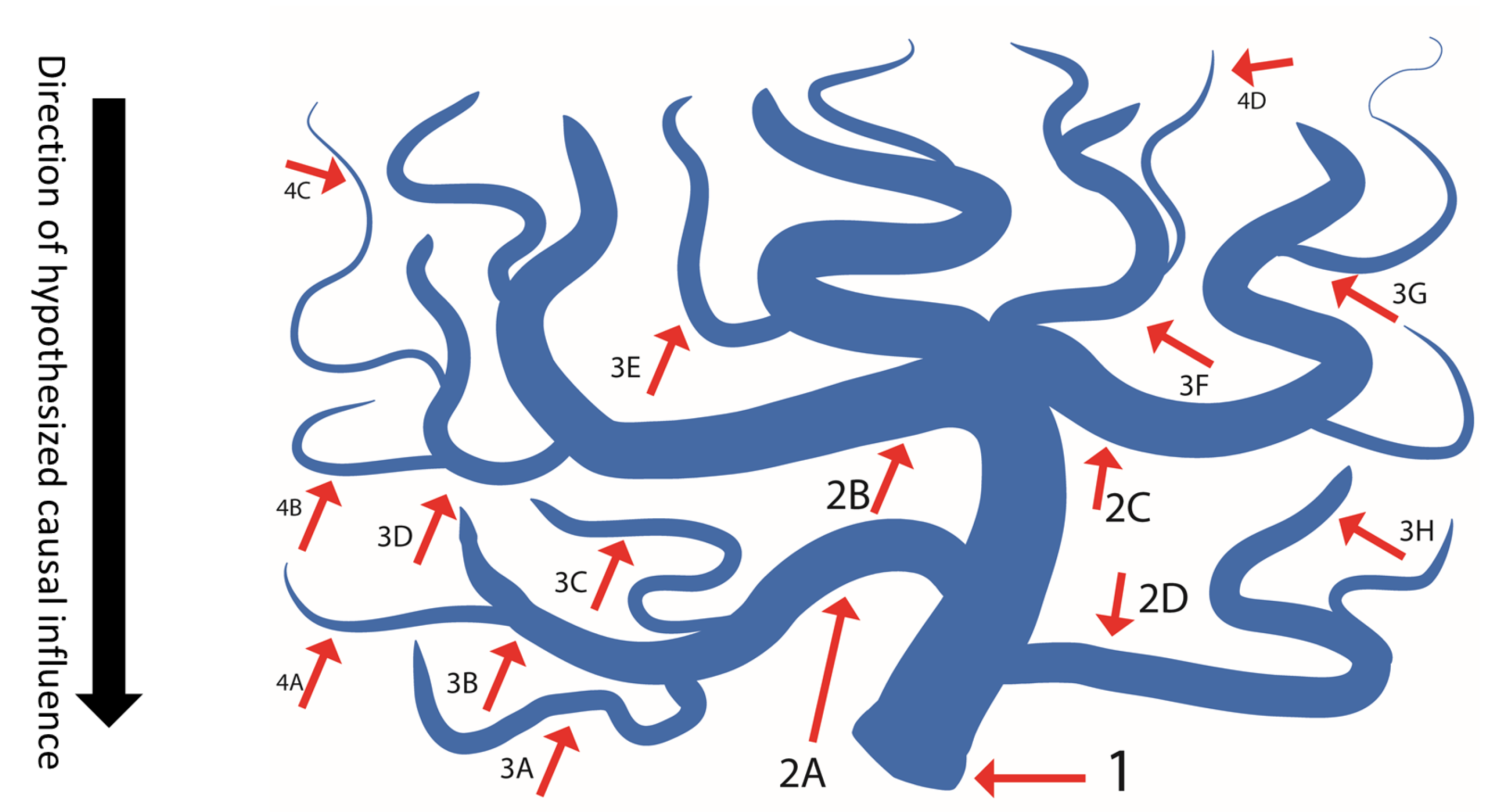
A watershed model of psychopathology (adapted from Cannon & Keller, 2006). Point “1” represents the most complex phenotype, such as schizophrenia (or fluid intelligence for our purposes). Points “2a-2d” represent endophenotypes, such as lower-level behavioural consequences; points “3a-3g” represent the neural antecedents of those behavioural phenotypes. Points 4a-4d represent hypothetical genetic influences (not measured here)

This model has a variety of conceptual benefits, including the fact that it naturally accommodates the constellations of antecedent causes that can contribute, independently, to some aggregate behavioural phenotype. This proposed causal heterogeneity explains the relative lack of success of directly mapping phenotypes onto genetic markers: Given that there is inherent multiplicity in the phenotype, the path from genetic causes to phenotypic outcomes will be noisy, so large samples will be needed (see Ripke et al., 2014 for an illustration of the striking increase in variance explained once sample sizes are large enough to estimate many small effects).

Cannon and Keller (2006, p. 274) derive from their model various empirical and conceptual predictions. These include that endophenotypes (intermediate causes) should be heritable, they should be associated with causes rather than effects, and numerous endophenotypes should affect a given construct. The model therefore predicts that a more efficient way of studying genetic causes is to focus on the endophenotypes of a disorder first, and then examine the genetic antecedents of those endophenotypes located further ‘upstream’. Moreover, endophenotypes are expected to vary continuously in the population and they should affect multiple disorders. We here adopt the watershed model to explain the relationship between fluid intelligence, processing speed and white matter by translating the watershed model from conceptual tool into a testable statistical model.

In our representation, FI is the ‘mouth’ of the river, influenced by upstream endophenotypes of PS and WM. This implies a hierarchical relationship, such that greater WM organisation (3a-3h) affects PS (2a-2d), which in turn affects fluid intelligence (1). If we examine the predictions by Cannon and Keller described above, we can see that both the phenotype and the proposed endophenotypes are highly heritable-FI (Deary et al., 2010), PS (e.g. Vernon, 1989) and whole-brain FA (Chiang et al., 2009)-yet there has been a notable lack of success in establishing replicable genetic markers for FI(Chabris et al., 2012). Recent work has shown how, under certain circumstances, reductionist hypotheses like that of Cannon and Keller can be translated to formal statistical models, such as structural equation models (SEM) of covariance patterns (Kievit, 2014; Kievit et al., 2011a, 2011b; Salthouse, 2011). Below, we show how the multiple predictions of the watershed model can be translated into statistical tests within a structural equation modelling (SEM) framework.

### 1.3.1 Greater upstream statistical dimensionality

As one can see in *Figure 1*, upstream ‘tributaries’ represent partially independent influences on the phenotype. This means that if we move up the tributaries of the river and examine the statistical dimensionality of the variables at each level, we would expect the dimensionality of the covariance pattern between all variables at that level to increase. Although they likely share some environmental or genetic influences and so will be correlated to some degree, we expect that the upstream effects cannot be fully captured by a single summary statistic. More importantly, we expect these upstream effects to be partially independent, such that any one intermediate endophenotype in isolation will do worse than a broader set in terms of predicting the downstream outcome.

### 1.3.2 Multiple realizability

As we have seen above, the watershed model suggests that seemingly unitary constructs are nonetheless likely to have multidimensional antecedent causes. In other words, a single behavioural dimension such as intelligence is likely to have multiple neural determinants; a type of between-individual *degeneracy* (Friston and Price, 2003). There is increasing support for such a many-to-one brain-behaviour mapping. For example, recent evidence suggests that differences in emotional states are better seen as a broad network of regions showing a different activation profile, rather than activity in individual regions in isolation mapping onto individual emotional states (Lindquist et al., 2012). Similarly, many concurrent and partially independent neural properties determine individual differences in broad cognitive skills such as general intelligence (Kievit et al., 2012; Ritchie et al., 2015b). In a SEM framework, this prediction means that variability in each endophenotype will make partially independent contributions to variability in the phenotype, in line with a so-called *MIMIC* model (Multiple Indicators, Multiple Causes; see Jöreskog and Goldberger, 1975; Kievit et al., 2012).

### 1.3.3 Hierarchical dependence

A defining characteristic of the watershed model is *hierarchical dependence.* That is, the influence of upstream causes are presumed to ‘flow through’ lower levels (endophenotypes). Statistically, this implies that there should be no residual, or direct, relationships between levels separated by a purported endophenotype. In the present context, we hypothesize that the influence of white matter is indirect, namely through processing speed. In the SEM formalization of this hypothesis below, any direct paths between WM and FI will be a source of model misfit. Taken together, it is possible to capture all these statistical predictions in a single structural equation model. This model is a hierarchical version of the MIMIC model (shown graphically in Figure 6). This model assumes that a latent variable (here FI) represents the phenotypic endpoint. The unidimensionality of this phenotype is tested by fitting a confirmatory factor model to the various behavioural measures available (four sub-scores of the Cattell test in the present data). At the second level, we hypothesize that various measures of PS: a) cannot be captured by a single factor, b) provide partially independent predictions of the fluid intelligence, and c) the latent variable of FI ‘shields off’ all direct effects of speed measures on the observed Cattell scores. Likewise, the WM tracts should have partially independent influences on the PS variables, but there should be no direct paths to FI that would explain away the relation between PS and FI. The statistical predictions described above can either be tested as part of the full model such that violations will lead to model misfit, or by explicit testing of individual predictions using model comparison. We will fit a MIMIC model in stages, so as to build up to the full the watershed model, and examine whether the predictions at the various stages described above are supported by our data.

## 2 Method and Experimental Procedures

### 2.1 Sample

A healthy, population-derived sample was collected as part of Phase 2 (“700”) of the Cambridge Centre for Ageing and Neuroscience (Cam-CAN), described in more detail in (Shafto et al., 2014). Exclusion criteria included low Mini Mental State Exam (MMSE) (24 or lower), poor hearing, poor vision, poor English (non-native or non-bilingual English speakers), self-reported substance abuse and current serious health conditions. Prior to analysis, we defined outliers as values for any variable that were more than 4 standard deviations from the mean (0.27% of all values) and included all participants with scores on all variables. The final sample contained 555 people, 274 female, age approximately uniformly distributed across the age range 18-87 (M=53.96). Table 1 contains descriptives of the sample in terms of sex, basic cognitive function and health factors. Note that, because our sample is cross-sectional, the findings relate to individual differences that are compatible with the watershed model, which may be, but are not necessarily, the same as age-related changes within an individual (e.g. Hofer & Sliwinski, 2001). A subset of these data have been reported in (Kievit et al., 2014). The covariance matrix is in the Supplementary Table 1. The raw data and analysis code are available upon signing a data sharing request form (see http://www.mrc- cbu.cam.ac.uk/datasets/camcan/ for more detail). As our sample included older participants, we report prevalence of common cardiovascular conditions in Table 1.

**Table 1:** Sample descriptives of the N=555 sample.

| Variable | Mean (SD) |
| --- | --- |
| Age | 53.96 (18.27) |
| MMSE | 28.85 (1.37) |
|  | % |
| Sex (female) | 49.29 |
| Hypertension diagnosis | 18.55 |
| Diabetes diagnosis | 3.96 |
| Myocardial infarct | 0.54 |
| Miscellaneous cardiovascular<br>(Cardiac arrhythmia/palpitations/ irregular heartbeat) | 7.21% |

### 2.2 Processing speed

Processing speed reaction times on three different cued-response tasks: simple response time (SRT), choice response time (CRT) and audio-visual cued response time (AV). These tasks differed in their demand characteristics, such as the nature of the cue and the predictability of the stimulus, and so may tap different aspects of PS. The AV task in particular differed from the SRT and CRT in that participants were not explicitly instructed to respond as quickly as possible, so this variable captures the natural response time in the absence of time pressure. For procedural details, see Figure 2A–C and Shafto et al., 2014: p. 6 (for SRT and CRT) and p. 16 (for AV). We include the mean and standard deviation for all three tasks, leading to a total of 6 measures of. All six variables were scaled to a standard normal distribution, log transformed and inverted (1/RT), so that higher scores reflect speedier and more consistent responses respectively (henceforth: SRT_speed_, CRT_speed_, AV_speed_, SRT_cons_, CRT_cons_ and AV_cons_).

**Figure 2:**
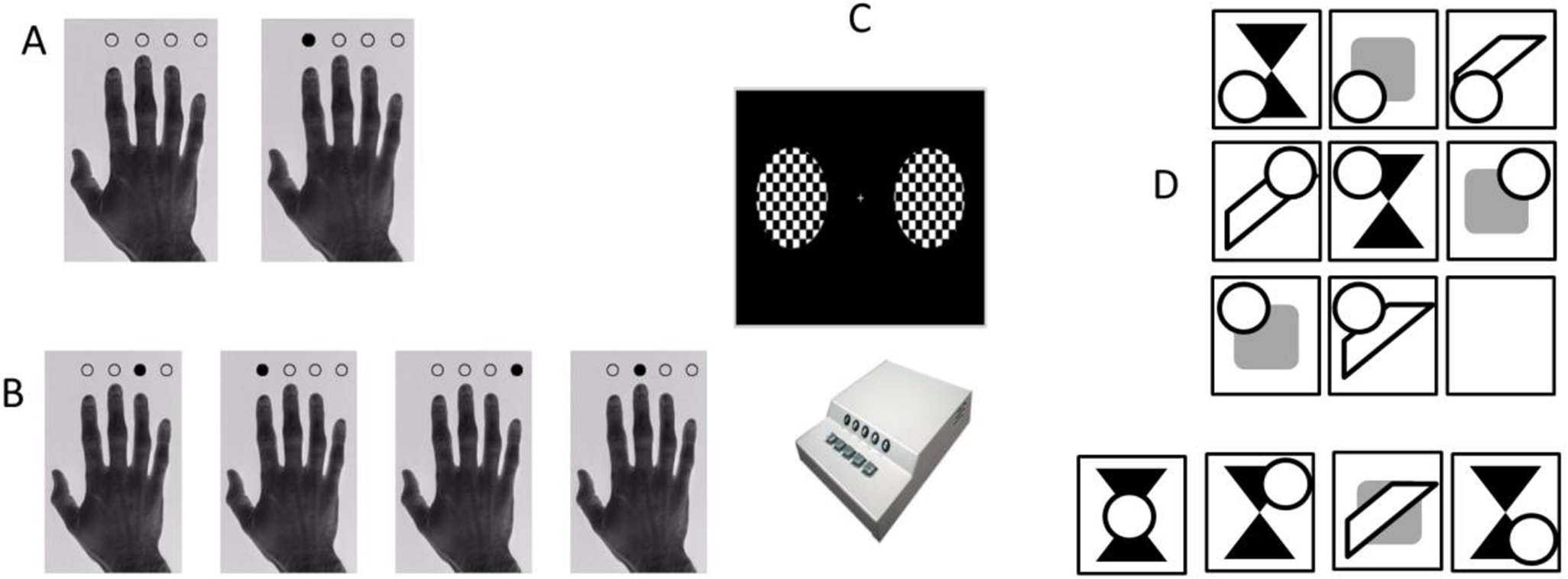
Behavioural measurements. Simple RT (a), choice RT (b), AudioVisual RT (C) and Cattell (d) (fictional example item shown).

### 2.3 Fluid intelligence

FI was measured using the Cattell's Culture Fair, Scale 2, Form A (Cattell, 1971), administered according to the standard protocol. This is a pen-and-paper test, consisting of four subtests with different types of abstract reasoning tasks, including series completion, classification, matrices and conditions. These four subtests each yield a sum-score representing the total number of correct responses which was scaled to a standard normal distribution prior to further analysis. Figure 2d shows an example test item.

### 2.4 White matter Organisation

In order to assess how different white matter tracts contribute to different cognitive functions, we computed mean Fractional Anisotropy (FA) values in various regions of interest (ROIs). Nonetheless, it should be noted that FA is a complex measure, and the relationship between FA and white-matter health is not yet fully understood (Jones et al., 2013; Wandell, 2016). There are a number of alternative measures that can be derived from diffusion-weighted MR images, which avoid the simplified single-tensor model, but the physiological validity of these is still under development (Tournier et al., 2011). A model focused on the contribution of the constituent physiological characteristics of white matter would be ideal for future applications of the watershed model, e.g. to test the complementary roles of axonal structure vs. myelin fraction (Caspers et al., 2015; Seehaus et al., 2015). Because our model does not make specific predictions for specific cellular constituents (e.g., water fraction, axonal diameter, myelin density), we favour the use of a simple tensor measure of diffusion organization, fractional anisotropy (FA). Most importantly, the vast majority of studies of white matter in healthy aging have used FA, and FA has been shown to be a comparatively reliable metric (Fox et al., 2012). For more detail on the white matter pipeline, see Appendix A. We computed mean FA for ten tracts as defined by the Johns Hopkins University white matter tractography atlas (Hua et al., 2008): The Uncinate fasciculus (UNC), superior longitudinal fasciculus (SLF), inferior Fronto-occipital fasciculus (IFOF), anterior thalamic radiations (ATR), forceps minor (FMin), forceps major (FMaj), cerebrospinal tract (CST), the inferior longitudinal fasciculus (ILF), ventral cingulate gyrus (CINGHipp) and the dorsal cingulate gyrus (CING) – see Figure 5A.

### 2.5 SEM

All models were fit using the package Lavaan (Rosseel, 2012) in R (R Development Core Team, 2016). Prior to model fitting, variables were scaled to a standard normal distribution and log transformed where necessary to increase normality. All models were fit using Maximum Likelihood Estimation (ML) using robust standard errors and report overall model fit assessed with the Satorra-Bentler scaled test statistic. Model fit was also assessed with the chi-square test, RMSEA and its confidence interval, the Comparative Fit index and the standardized root mean squared residuals (Schermelleh-engel et al., 2003). We define good fit as follows: RMSEA<0.05 (acceptable: 0.05–0.08), CFI>0.97 (acceptable: 0.95–0.97) and SRMR < 0.05 (acceptable: 0.05–0.10) and report the Satorra-Bentler scaling factor for each model. Models are compared using a chi-square test when nested and using the AIC in other cases.

## 3. Results

To examine the predictions of the watershed model, we will build up the full model, starting at the ‘top’. First, we fit our measurement model, namely relating the latent variable FI to the four scores on the Cattell subtests. This model fit the data well: χ^2^=4.372 (N=555), df=2, p=0.112, RMSEA=0.046 [0.000 0.106], CFI=.996, SRMR = 0.011, Satorra-Bentler scaling factor=1.018, suggesting that performance could be captured by a single dimension, as predicted by the watershed model. As expected, scores on the latent variable showed steep age-related decline. A linear regression explained 43.97% of the variance (*N*=555, *F*(1,553)=435.8, p < 0.0001, adjusted R^2=43.97%) and a second-order polynomial explained (adjusted) 46.82% of the variance (*N*=555, *F*(2,552)=244.8, p < 0.0001), with the AIC slightly favouring the polynomial (steeper decline in later life, AIC_linear_=1163.74, AIC_poly_=1135.82). Furthermore, a Breusch-Pagan test showed that residuals increased slightly with age, suggesting greater inter-individual variability in later life (BP=18.658, df=2, p<0.0001). Model fit and age-related differences are shown in Figure 3.

**Figure 3.**
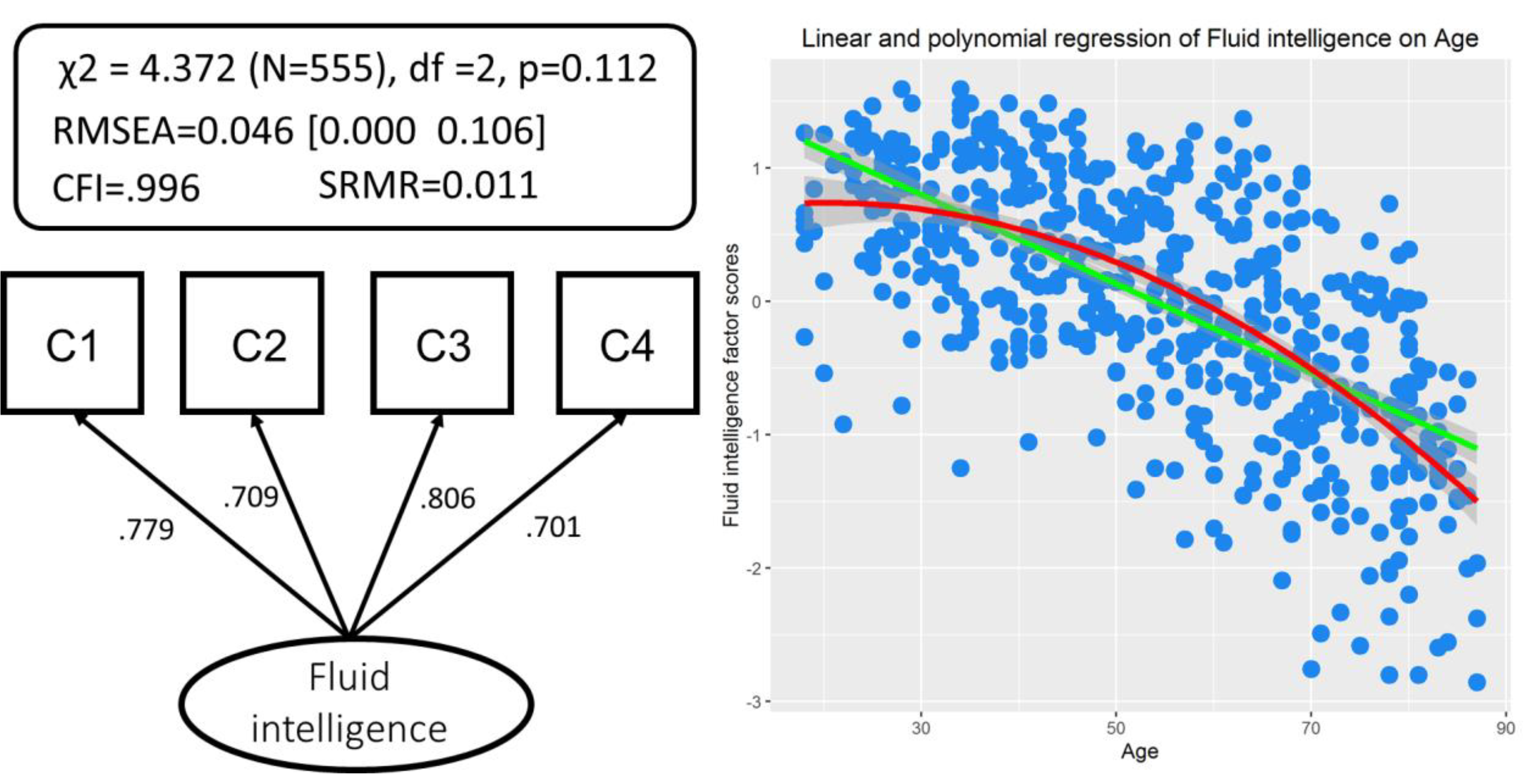
A single-factor confirmatory factor analysis for Cattell fits data well (left). Linear and polynomial fit of age-related differences in fluid intelligence (right).

In order to establish the relationship between PS and FI, we first examined the dimensionality of the PS measures. Since the watershed model suggests that variables more ‘upstream’ may be partially independent, we first tested whether a single unidimensional model fit the six PS measures (see also Babcock, Laguna, & Roesch, 1997). A single factor model (Supplementary Figure 1A) to the six PS measures showed poor fit χ^2^=696.67, df=9, p < 0.0001, RMSEA=0.371 [0.349 0.394], CFI=0.549, SRMR=0.162, Satorra-Bentler scaling factor=1.08. We then examined whether a model with two latent variables (Supplementary Figure 1B), one for speed (measured by SRT_speed_, CRT_speed_, AV_speed_) and one for consistency (SRT_cons_, CRT_cons_ and AV_cons_) would fit better. This model also fit poorly: χ^2^=663.940, df=8, p < 0.0001, RMSEA=0.384 [0.361 0.408], CFI=0.57, SRMR=0.158, Satorra-Bentler scaling factor=1.13 as did an alternative model (Supplementary Figure 1C) with a latent factor for each task (SRT, CRT and AV) χ^2^=122.691, df=6, p < 0.0001, RMSEA=0.187 [0.160 0.216], CFI=0.923, SRMR=0.048, Satorra-Bentler scaling factor=1.084). This suggests that our measures of PS cannot be reduced to a single dimension. However, a more crucial question in the context of the watershed model is whether the upstream antecedents will make partially independent contributions to fluid intelligence. To test this hypothesis, we fit the simple MIMIC model shown in Figure 4.

**Figure 4.**
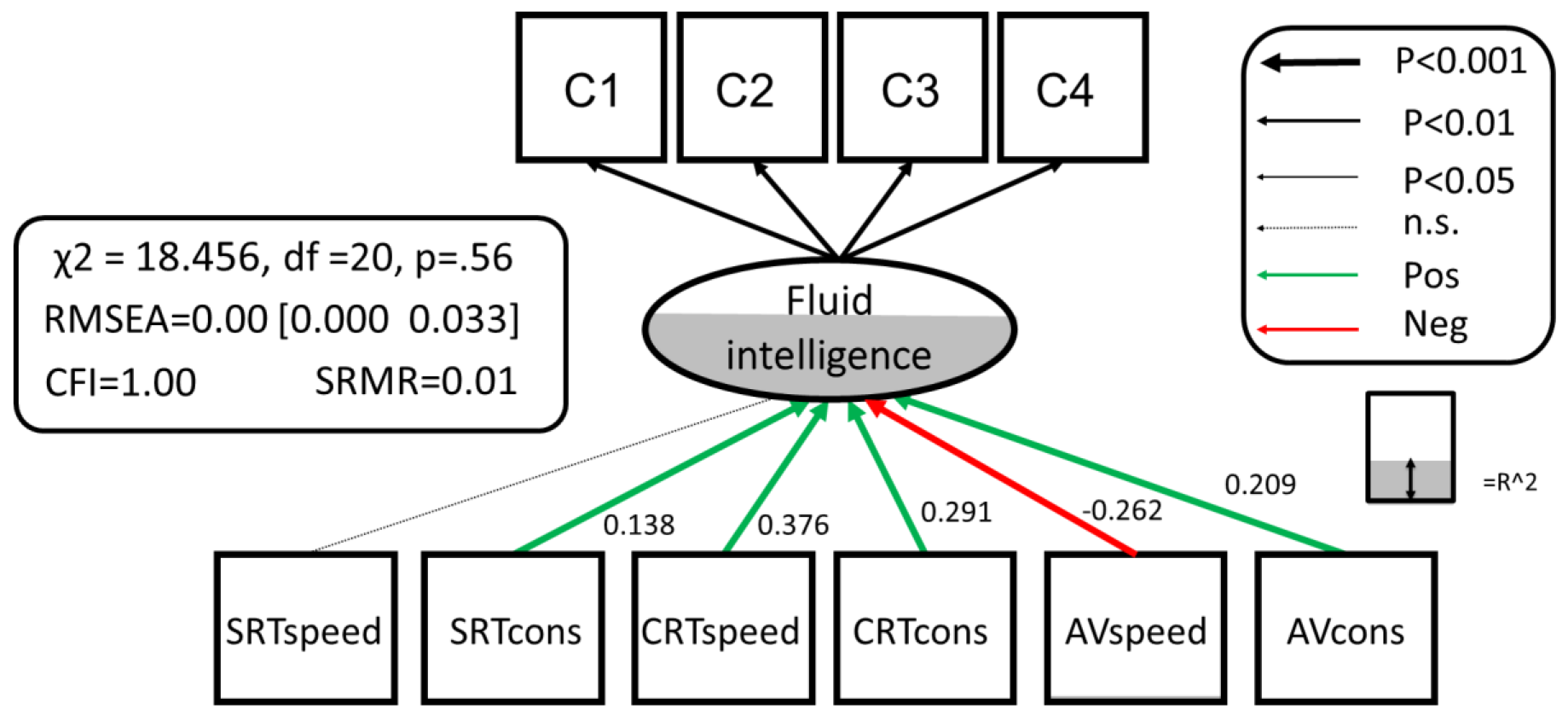
MIMIC model for Processing Speed and Fluid intelligence. Below age-related trajectories for each processing speed measure ranging from strong CRT_speed_, r=−0.64) to absent (AV_speed_, r=0.03, n.s.). Residual covariances between PS variables are allowed but not shown for simplicity.

This showed excellent fit to the data: χ^2^=18.456, df=20, P=.56, RMSEA=0.000 [0.000 0.033], CFI=1.00, SRMR=0.010, Satorra-Bentler scaling factor=1.034. Most strikingly, five out of the six PS variables (all but SRT_speed_) predicted unique variance in fluid intelligence. Next, we compared this model, with all PS to FI pathways estimated freely, to more parsimonious competitors. First, a simple model where all PS to FI pathways were set to 0 fit considerably worse than a model with PSto FI pathways estimated freely (χ^2^Δ=331.72, dfΔ=6, p <.0001). Second, amodel where all PS to FI pathways were constrained to be equal (Supplementary Figure 1D) again fit worse than the full model (χ^2^Δ=175.5, dfΔ=5, p <.0001), suggesting specificity of the pathways in line with the watershed model. Finally, we tested whether only estimating the strongest pathway (CRT-speed) and constraining the other pathways to 0 might suffice (Supplementary Figure 1E). This model too showed worse fit than estimating all six pathways freely (χ^2^Δ=69.668, dfΔ=5, p <.0001). Together these comparisons show that the relationship between processing speed is many-to-one, and cannot be captured fully by a single pathway, supporting the suggestion by Salthouse (2000) that different indicators of PS may reflect ‘somewhat distinct processes’ (p. 41). Figure 4 shows the partially independent contributions of the PS measures to FI that together explain 58.6% of the variance in FI. Perhaps surprisingly, AV_speed_ had a modest *negative* path, suggesting that those with higher FI scores are those who had fast response speed when instructed to respond quickly, but slower response speed when not so instructed. These findings suggest that different tasks and instructions can tap into distinct underlying processes, and that individual differences in these processes combine to explain a considerable portion of individual differences in FI. In other words, the results suggest that mental speed is multifaceted and that different elements play complementary roles in supporting higher cognitive abilities (Schubert et al., 2015).

Finally, the ‘lowest’ layer of our model pertains to the structural organization of WM as evidenced by the isotropy of water diffusion in major WM ROIs. The covariance of WM structure across individuals is informative because its dimensionality can suggest possible mechanisms driving individual differences, which has been a subject of contention in the literature (e.g. Kievit et al., 2014; Lövdén, Laukka, et al., 2013; Ritchie, Bastin, et al., 2015). We fit SEMs to the mean FA of the ten, bilaterally averaged, WM tract ROIs (Figure 5a), which showed different sensitivities to age (Figure 5B). A single factor model showed poor fit (χ^2^ = 422.182, df = 35, p < 0.0001, RMSEA = 0.141 [0.130 0.153], CFI=0.819, SRMR =0.069, Satorra-Bentler scaling factor=1.138). The poor fit of the single-factor model was not driven by differential ageing of the tracts (Figure 5B): We refit the model to the age-corrected residuals of the 10 tracts, but this also fit the data very poorly (χ^2^ = 454.917, df = 35, P<0.0001, RMSEA =0.147 [0.136 0.158], CFI = .747, SRMR =0.079, Satorra-Bentler scaling factor=1.132). Inspection of the modification indices showed no simple modifications that would show better fit, suggesting covariance in white matter organization in our sample cannot be reduced to a single dimension. We now move to fitting the full model, including all levels simultaneously.

**Figure 5.**
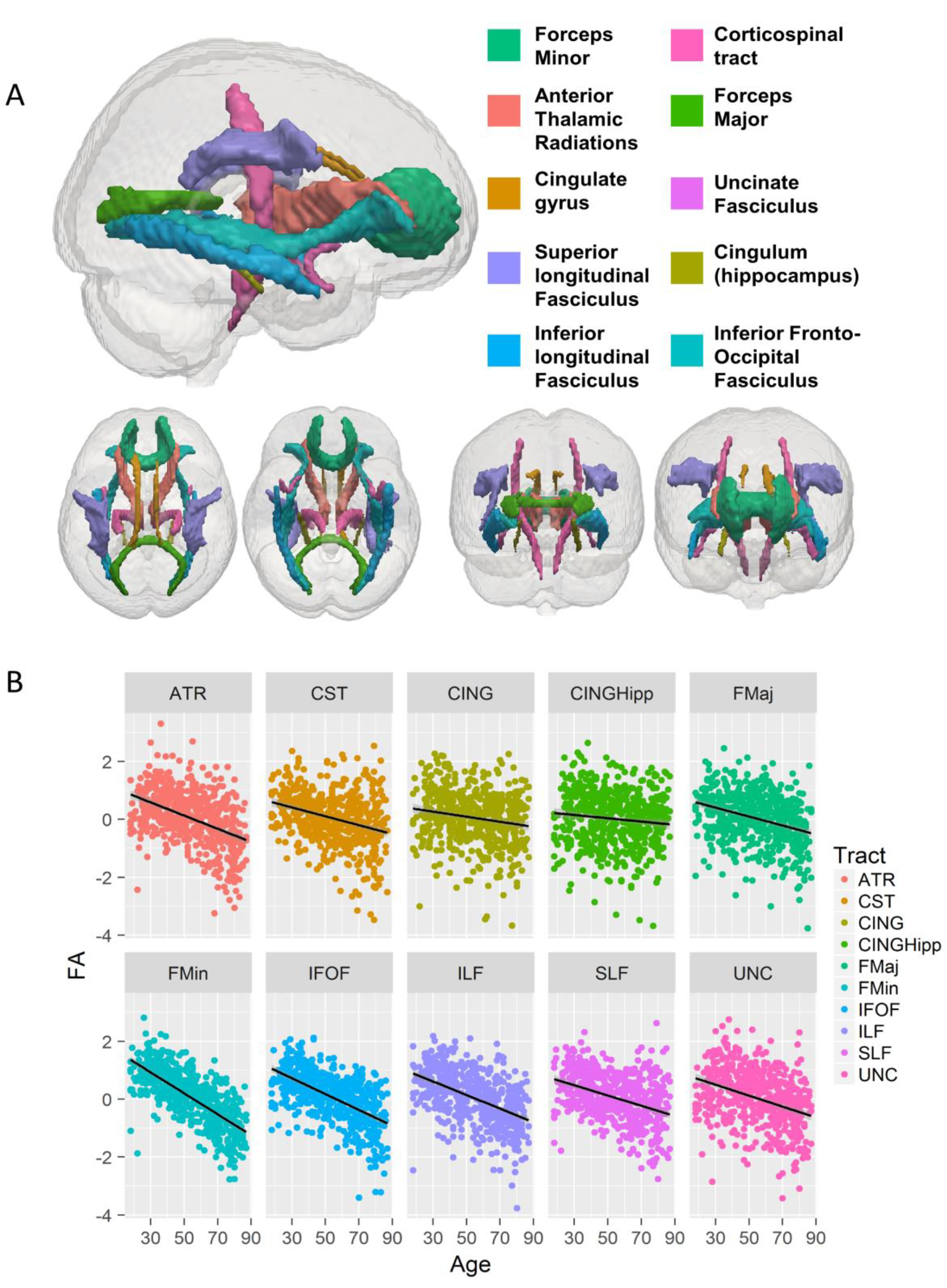
A) All ten white matter tracts used in our analysis, based on the JHU Atlas. B) Differential ageing of the ten tracts, correlations ranging from −0.71 (Forceps Minor) to −.10 (Ventral Cingulum, or CINGHipp).

### 3.1 Full model

So far, the findings are in line with the predictions of the watershed model. We then fit the full watershed model (Figure 6), integrating fluid intelligence, PS and FA in a set of major WM tracts. By doing so, we can simultaneously test the hierarchical structure and many-to-one mapping imposed by our conceptual framework. In this model, we allow for residual covariances *within,* but not *between* levels. This model captures the assumption that all influence that WM tracts have on FI should go through the PS level (i.e., any residual covariance between WMI and the latent variable of FI, or any of the subtests, would be a source of misfit). Finally, any influence of PS on the Cattell subtests should go via the latent variable of FI. The full model, as shown in Figure 6 fits the data very well: χ^2^ = 103.201, df = 60, p<0.001, RMSEA =0.036 [0.024 0.047], CFI =.986, SRMR =0.034, Satorra-Bentler scaling factor=1.033. This suggests that the observed covariance pattern in our data is compatible with the statistical constraints imposed by the watershed model. In other words, that the data are compatible with the hypotheses that the three explanatory levels stand in a hierarchical relationship, such that WM determines PS, which in turn determines FI. Given that FA is known to be affected by vascular health, we also added four binary cardiovascular/general health conditions in Table 1 as covariates (affecting white matter FA). This had little effect on model fit, χ^2^=156.280, df = 100, P<0.001, RMSEA=0.032 [0.022 0.041], CFI=.989, SRMR=0.039, Satorra-Bentler scaling factor=1.122, suggesting that the results from the full model did not simply reflect unmodelled differences in vascular health.

A key prediction from the watershed model is that the relationship between upstream measures and downstream consequences is many-to-one. We can test this hypothesis by investigating, as we did previously when relating PS to FI, whether more parsimonious accounts of the relationship between WM and PS show better fit. First, we found that a model with all WM to PS pathways constrained to 0 showed poor fit (χ^2^Δ = 311.8, df Δ = 60, p<0.001). Second, we tested a ‘strongest path only model’, estimating the Forceps Minor pathway but fixing all others to 0 (Supplementary Figure 2A). This model too showed worse fit χ^2^Δ = 80.31, df A = 54, p<0.05. Next, we tested a ‘white matter tract specificity’ model. Here we allow the effect of white matter to vary between tracts, but to be equal across the six processing speed measures (Supplementary Figure 2B). This model also fit worse, χ^2^Δ=142.76, df Δ =40, p<0.001). Finally, we tested a ‘processing speed specific’ model, in which the effects were allowed to differ across processing speed measures, but constrained to be equal for each tract (Supplementary Figure 2C). This model again showed poorer fit than the full model (χ^2^Δ=142.45, df Δ=54, p<0.001).

**Figure 6:**
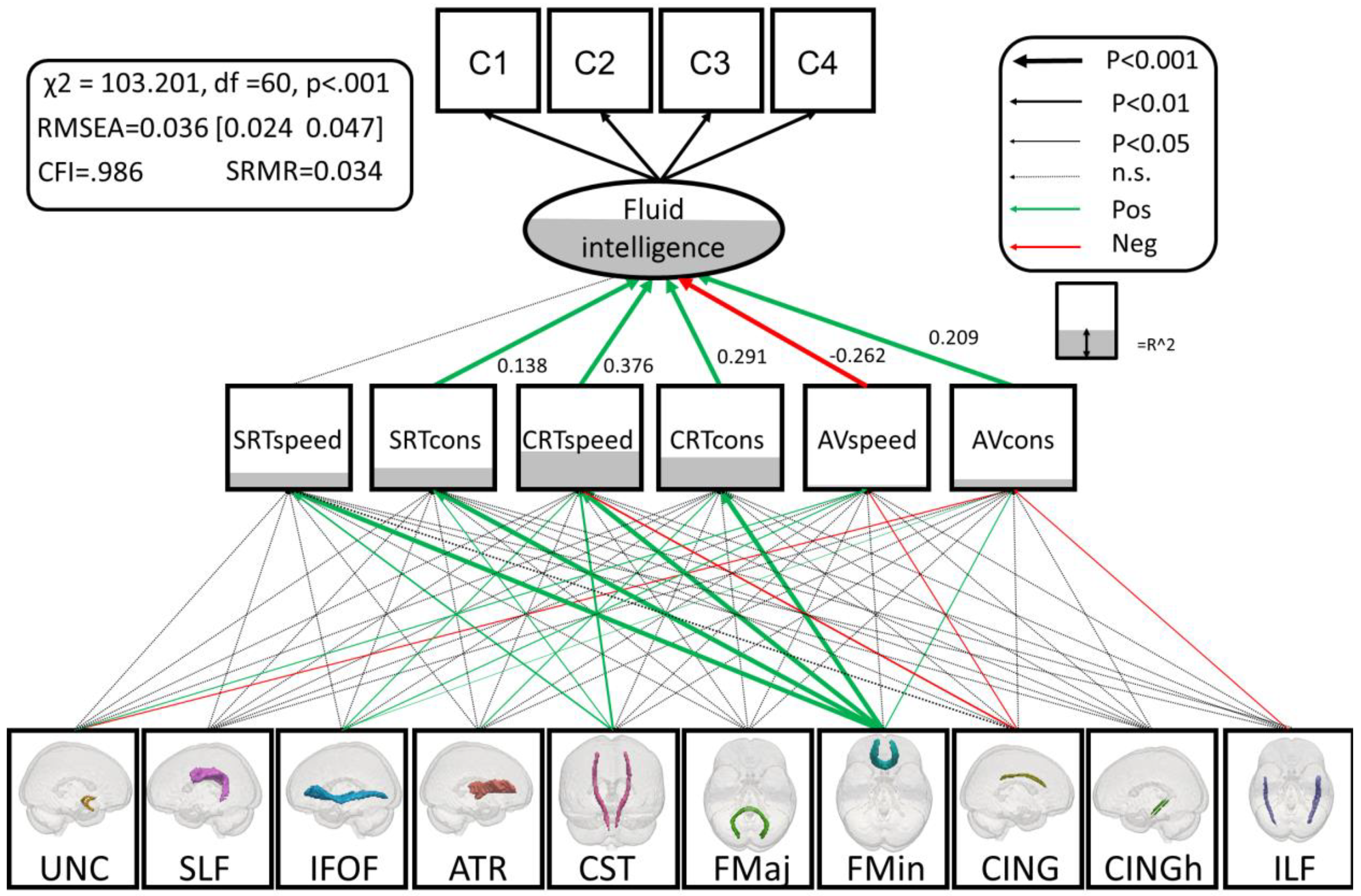
Full watershed model. Significant parameters are shown in green and red, R-squared is represented as the degree of shading of the variables. Residual covariances between processing speed variables and white matter tracts are allowed, but not shown for simplicity. See Supplementary Table 1 for the full covariance matrix, and Supplementary Table 2 for the unstandardized parameter estimates and se's.

Finally, to establish that the fit of the full model was not merely a consequence of some more general property of the covariance matrix, we performed control analyses to test the nature of the hierarchical relationship. Firstly, we *inverted* the two lower levels, such that PS affected WM organisation which in turn directly affected FI (again allowing for residual covariances between all WM tracts, but precluding direct influence between PS and fluid intelligence). The original watershed model fit the data considerably better (AIC_diff_=224.28). Secondly, because there are more WM variables than PS variables (which might affect model comparisons), we exhaustively compared all 210 combinations of 6 tracts (to match the number of PS variables) to an inverted model with the same subset of tracts. In every model comparison, the watershed model outperformed the inverted model (AIC_diff_ ranged from 141.76 to 264.85 in favour of the watershed model). Again, including cardiovascular factors as WM covariates had little effect, with the watershed model being preferred to the inverted model in every combination (AIC_diff_ ranged from 150.41 to 253.03 in favour of the watershed model).

One notable observation was that the strongest prediction of PS was WM organisation in the Forceps Minor (also known as the anterior forceps, which passes through the genu of the corpus callosum). This variable explained significant amounts of variance in five of the six response time measures. Moreover, this relationship was strongest for the most ‘executive’ of PS variables, the speed and consistency of the Choice RT task, in line with previous findings that suggest an important role for prefrontal WM in such tasks (Davis et al., 2009). In fact, inspection of the modification indices suggested a possible residual, direct pathway from the Forceps Minor to FI, and adding this pathway did indeed lead to improved fit (χ^2^Δ = 36.297, dfΔ=1, p<0.001). This suggests the possibility of a cognitive endophenotype between white matter and FI that was not captured by our PS measures. A likely candidate for this endophenotype to explore in future work is working memory capacity (e.g. Fry & Hale, 1996). Importantly however, adding this direct pathway did not affect (e.g. render non-significant) the existing PS to FI paths, supporting an independent pathway rather than a violation of the proposed hierarchy.

The second strongest effect was from the cortico-spinal tract, which affected three speed measures above and beyond the variance already captured by Forceps Minor (see also Duering et al., 2013, and Lövdén et al., 2014). Together, these findings provide significant support for the watershed model: white matter, processing speed and fluid intelligence stand in a hierarchical, many-to-one relationship that requires measuring a broad spectrum of variables at each explanatory level.

## 4. Discussion

In our population-based, age-heterogeneous sample, we found strong evidence for a hierarchical relationship between fluid intelligence, processing speed and white matter. Specifically, individual differences in WM anatomy predicted individual differences in processing speed, which in turn predicted over 58% of the variance in fluid intelligence scores. This model performed significantly better than control models that inverted these explanatory levels, or models that imposed equal or summary effects between levels. This watershed model is based on theoretical considerations, is in line with a wealth of empirical evidence, and provides an overarching framework for modelling causal relationships. All statistical predictions derived from the watershed model were supported by our data. First, our outcome measure (fluid reasoning) fit a single factor model, whereas no low dimensional models fit for either PS or WM. Second, there was evidence for a many-to-one mapping, such that multiple individual variables at the processing speed and WM levels explained unique variance in higher explanatory levels. Third, the overall model fit significantly better than various alternatives, providing evidence for hierarchical dependency. These findings have important implications for both the understanding of age-related declines in intelligence, and for the proscriptive ability of cognitive neuroscience to potentially inform successful interventions within aged communities. We expand on the findings and implications below.

Five out of the six processing speed measures predicted unique variance in fluid intelligence, supporting the hypothesis that processing speed is a multidimensional construct, and that these subtle aspects are important for understanding higher, abstract cognitive abilities. Similarly for WM, and in line with recent work (Lövdén et al., 2013a), we found that individual differences in WM are multidimensional, and that these different dimensions have partially independent predictions for processing speed. The strongest influence was that of the Forceps Minor, which predicted five distinct processing speed measures, with the Corticospinal Tract and the Inferior Fronto-Occipital Fasciculus also explaining considerable variance in PS measures. In contrast to views that aging represents a monolithic decline, the current findings support the idea that distinct brain regions and distinct cognitive abilities change in different ways, and that only models which strive to incorporate this multiplicity in their explanations of age-related decline can capture the entirety of age-related processes (Andrews-Hanna et al., 2007; Kievit et al., 2014; Lövdén et al., 2014; Tucker-Drob, 2011).

These findings may have implications for the design and implementation of cognitive training interventions. The watershed model suggests that transfer to other cognitive domains (see also (Taatgen, 2013) may only be achieved if the intervention is of sufficient length and intensity to affect the entire hierarchy of relationships, including the mapping of lower processing speed to higher cognitive processes. Such a suggestion is supported by work showing that white matter supports many distinct cognitive functions (Burzynska et al., 2013). For example, to truly improve fluid reasoning and not just observed scores (Hayes et al., 2015), cognitive training would have to be of such duration that lower levels (such as processing speed and WM) are also measurably affected (Keller and Just, 2009; Scholz et al., 2009), which can then generalize to other domains (Lövdén et al., 2010a). Behavioural evidence for such a pattern was found by Edwards et al. (2002), who showed transfer of speed of processing training to multiple cognitive domains in older adults. Additional evidence for this possibility comes from study by Schmiedek et al. (2010) who observed modest cognitive transfer as well as white matter microstructural change (Lövdén et al., 2010b) after a high intensity (100 day) cognitive training. Notably, the positive effects remained for up to two years (Schmiedek et al., 2014).

In addition to the empirical findings reported here, there are methodological advantages to implementing this model. By visualizing the full model, including statistical quantification of the strengths of association such as R^2^, it immediately emphasizes not just which ties are strong and well-established, but also shows where our knowledge is lacking. For example, not all variance can be explained in either fluid reasoning or processing speed (see also Rabbitt et al., 2007b), suggesting we need to explore other metrics of processing speed (e.g. inspection time, or digit-symbol substitution), additional cognitive determinants (e.g. working memory; Engle, Tuholski, Laughlin, & Conway, 1999; Fry & Hale, 1996), and additional neural markers such as prefrontal activity (e.g. Christoff et al., 2001), and structure (Waltz et al., 1999; Woolgar et al., 2010), grey matter indices (Kievit et al., 2014; Stuss et al., 2003) or additional WM metrics such as MD and AD (Tamnes et al., 2012) to obtain a more complete picture.

One limitation of the model as implemented here is that our sample is cross-sectional, not longitudinal. This means that although we can model the extent to which *individual differences* are in line with the watershed model, but we cannot make claims concerning intra-individual changes over the lifespan (Raz & Lindenberger, 2011; Salthouse, 2011, Kievit et al., 2013). To truly get at the developmental dynamics, it would be necessary to follow people over time, most crucially during the critical periods of adolescence and later-life aging. The watershed model would predict that, at sufficient temporal resolution, developmental changes in white matter organisation would precede changes in processing speed, which in turn precede changes in fluid reasoning. A second limitation is the selection of tasks. Although those we included cover four domains of fluid reasoning (series completions, odd-one-out, matrices and topology), they are all subtests of a single test. Ideally, a model should include additional fluid reasoning tasks (such as Raven's Matrices) to capture a broader spectrum of reasoning abilities. Similarly, all our processing speed measures focus on response time, where a broader spectrum of tasks tapping processing speed (e.g. inspection time or digit-symbol substation or diffusion model parameters, cf. Deary & Ritchie, 2016; Yang et al., 2014) would allow for even more detailed investigation of the key hypotheses tested here, as would expanding the range of white matter metrics to include other measures of diffusivity and measures of magnetisation transfer (e.g. Penke et al., 2012;Yang et al., 2014).

In summary, the watershed model provides a powerful conceptual framework that organizes our knowledge and generates testable models of the expected covariance patterns within and across individuals. A strength of the model is that it naturally accommodates extensions in both ‘directions’. For example, findings in this model could be integrated with the study of other, even broader phenotypes. One notable and important extension of the model would be the inclusion of long-and short-memory measures, or measures of attention, which assess both an important aspect of higher functioning and also are notable in their age-related loss. By integrating multiple hierarchical models the interrelationships between cognitive phenotypes may become clearer. More ambitiously, it may be possible to integrate a model fit in a sample like ours with a larger study of psychopathological disorders known to be associated with impairments to cognitive abilities similar to fluid intelligence (e.g. schizophrenia, Snitz et al., 2006).

Although we here do not include the ‘lowest’ level of the watershed model, namely genetic effects, recent evidence from two large neuroimaging and genetic samples shows striking convergence with the predictions that follow from the watershed model with respect to white matter and processing speed. Kochunov et al. (2016) use quantitative genetic models to show that, in two independent samples (N=145 and N=481), ‘Quantitative genetic analysis demonstrated a significant degree to which common genes influenced joint variation in FA and brain processing speed.’ (p.190), and conclude that ‘specific genes influencing variance in FA values may also exert influence over the speed of cognitive information processing’ (p. 19). This overlap is precisely what one would expect based on the conceptual framework presented here.

The advent of larger, multimodal neuroimaging cohorts will allow us to integrate previously isolated empirical findings into larger explanatory models, thereby mapping the mechanistic pathways in increasing detail. Ultimately, mapping the full hierarchy from genotypes to phenotypes may provide novel insights into the cascade of developmental effects on complex cognitive abilities in health and disease.

## Author contributions

RAK conceived the project, analysed the data, and designed the SEM figures. MC and RNH developed the white matter analysis pipeline. SWD and RAK made the white matter figures. RAK, SWD, JG and RNH wrote the paper. All authors contributed to the final written product. This article was written in the absence of any competing financial interests.

## Disclosures and acknowledgements

The Cambridge Centre for Ageing and Neuroscience (Cam-CAN) research was supported by the Biotechnology and Biological Sciences Research Council (grant number BB/H008217/1). We are grateful to the Cam-CAN respondents and their primary care teams in Cambridge for their participation in this study. We also thank colleagues at the MRC Cognition and Brain Sciences Unit MEG and MRI facilities for their assistance, specifically Tibor Auer, Rafael Henriques and Nitin Williams for their assistance in working on the white matter pipeline. RAK is supported by the Sir Henry Wellcome Trust (grant number 107392/Z/15/Z) and the by UK Medical Research Council Programme (MC-A060-5PR61). RNH was additionally supported by UK Medical Research Council Programme MC-A060-5PR10.

The Cam-CAN corporate author consists of the project principal personnel: Lorraine K Tyler, Carol Brayne, Edward T Bullmore, Andrew C Calder, Rhodri Cusack, Tim Dalgleish, John Duncan, Fiona E Matthews, William D Marslen-Wilson, James B Rowe, Meredith A Shafto; Research Associates: Karen Campbell, Teresa Cheung, Linda Geerligs, Anna McCarrey, Abdur Mustafa, Darren Price, David Samu, Jason R Taylor, Matthias Treder, Kamen Tsvetanov, Janna van Belle, Nitin Williams; Research Assistants: Lauren Bates, Tina Emery, Sharon Erzinçlioglu, Andrew Gadie, Sofia Gerbase, Stanimira Georgieva, Claire Hanley, Beth Parkin, David Troy; Research Interviewers: Jodie Allen, Gillian Amery, Liana Amunts, Anne Barcroft, Amanda Castle, Cheryl Dias, Jonathan Dowrick, Melissa Fair, Hayley Fisher, Anna Goulding, Adarsh Grewal, Geoff Hale, Andrew Hilton, Frances Johnson, Patricia Johnston, Thea Kavanagh-Williamson, Magdalena Kwasniewska, Alison McMinn, Kim Norman, Jessica Penrose, Fiona Roby, Diane Rowland, John Sargeant, Maggie Squire, Beth Stevens, Aldabra Stoddart, Cheryl Stone, Tracy Thompson, Ozlem Yazlik; and administrative staff: Dan Barnes, Marie Dixon, Jaya Hillman, Joanne Mitchell, Laura Villis. We thank Peter Lugtig for commenting on an earlier draft of Figure 6.

---

**Supplementary Table 1:** Full covariance matrix.

|  | age | cattell1 | cattell2 | cattell3 | cattell4 | SRTspeed | SRTcons | CRTspeed | CRTcons | AVspeed | AVcons | ATR | CST | CING | CINGHipp | FMaj | FMin | IFOF | ILF | SLF | UNC |
| --- | --- | --- | --- | --- | --- | --- | --- | --- | --- | --- | --- | --- | --- | --- | --- | --- | --- | --- | --- | --- | --- |
| age | 334.0514 | -10.2549 | -9.36614 | -10.8366 | -8.75185 | -6.32071 | -7.61586 | -11.3662 | -10.4928 | 0.614443 | -3.26972 | -7.61455 | -5.1145 | -2.92023 | -1.9006 | -5.16363 | -12.0934 | -9.14884 | -7.82955 | -5.91601 | -6.31008 |
| cattell1 | -10.2549 | 1.008516 | 0.525742 | 0.634577 | 0.552186 | 0.278308 | 0.34254 | 0.537375 | 0.528172 | -0.08106 | 0.122258 | 0.315279 | 0.17419 | 0.127317 | 0.096505 | 0.204971 | 0.419928 | 0.35834 | 0.293436 | 0.227084 | 0.259104 |
| cattell2 | -9.36614 | 0.525742 | 0.976242 | 0.564852 | 0.511104 | 0.248089 | 0.315461 | 0.502861 | 0.477232 | -0.05423 | 0.147565 | 0.294938 | 0.244512 | 0.135062 | 0.166594 | 0.122581 | 0.372896 | 0.311892 | 0.233897 | 0.240043 | 0.204627 |
| cattell3 | -10.8366 | 0.634577 | 0.564852 | 0.982144 | 0.538095 | 0.276586 | 0.371694 | 0.50264 | 0.512145 | -0.0495 | 0.161466 | 0.335607 | 0.215077 | 0.146657 | 0.105157 | 0.203382 | 0.46767 | 0.380358 | 0.310533 | 0.259658 | 0.308776 |
| cattell4 | -8.75185 | 0.552186 | 0.511104 | 0.538095 | 0.991546 | 0.242004 | 0.313125 | 0.4755 | 0.485654 | -0.05945 | 0.160406 | 0.309525 | 0.200693 | 0.15918 | 0.085073 | 0.152919 | 0.3997 | 0.316838 | 0.246517 | 0.22572 | 0.24521 |
| SRTspeed | -6.32071 | 0.278308 | 0.248089 | 0.276586 | 0.242004 | 0.916255 | 0.635715 | 0.526525 | 0.335243 | 0.28062 | 0.260556 | 0.214907 | 0.196732 | 0.142647 | 0.060728 | 0.133269 | 0.327474 | 0.258725 | 0.219337 | 0.205932 | 0.192378 |
| SRTcons | -7.61586 | 0.34254 | 0.315461 | 0.371694 | 0.313125 | 0.635715 | 0.910692 | 0.494334 | 0.426512 | 0.117797 | 0.226462 | 0.272787 | 0.191679 | 0.165075 | 0.065631 | 0.126915 | 0.365904 | 0.281843 | 0.238493 | 0.23145 | 0.208202 |
| CRTspeed | -11.3662 | 0.537375 | 0.502861 | 0.50264 | 0.4755 | 0.526525 | 0.494334 | 0.920998 | 0.744841 | 0.059523 | 0.225533 | 0.367568 | 0.230475 | 0.171736 | 0.086895 | 0.215377 | 0.493354 | 0.406352 | 0.320487 | 0.282275 | 0.282191 |
| CRTcons | -10.4928 | 0.528172 | 0.477232 | 0.512145 | 0.485654 | 0.335243 | 0.426512 | 0.744841 | 0.923351 | -0.00754 | 0.207365 | 0.361796 | 0.152616 | 0.203876 | 0.068002 | 0.184593 | 0.462903 | 0.378202 | 0.317319 | 0.252321 | 0.295929 |
| AVspeed | 0.614443 | -0.08106 | -0.05423 | -0.0495 | -0.05945 | 0.28062 | 0.117797 | 0.059523 | -0.00754 | 0.953141 | 0.650712 | -0.01655 | 0.054487 | 0.06025 | 0.014313 | 0.011221 | 0.015212 | 0.033963 | 0.024775 | 0.023844 | -0.02542 |
| AVcons | -3.26972 | 0.122258 | 0.147565 | 0.161466 | 0.160406 | 0.260556 | 0.226462 | 0.225533 | 0.207365 | 0.650712 | 0.873757 | 0.163322 | 0.12686 | 0.139782 | 0.049874 | 0.103675 | 0.204707 | 0.190048 | 0.12005 | 0.149826 | 0.1042 |
| ATR | -7.61455 | 0.315279 | 0.294938 | 0.335607 | 0.309525 | 0.214907 | 0.272787 | 0.367568 | 0.361796 | -0.01655 | 0.163322 | 0.890484 | 0.240021 | 0.400037 | 0.077276 | 0.208996 | 0.560993 | 0.570611 | 0.36261 | 0.424691 | 0.460745 |
| CST | -5.1145 | 0.17419 | 0.244512 | 0.215077 | 0.200693 | 0.196732 | 0.191679 | 0.230475 | 0.152616 | 0.054487 | 0.12686 | 0.240021 | 0.808179 | 0.244631 | 0.312649 | 0.124764 | 0.280685 | 0.317736 | 0.305891 | 0.372024 | 0.195309 |
| CING | -2.92023 | 0.127317 | 0.135062 | 0.146657 | 0.15918 | 0.142647 | 0.165075 | 0.171736 | 0.203876 | 0.06025 | 0.139782 | 0.400037 | 0.244631 | 0.850662 | 0.116428 | 0.093792 | 0.431222 | 0.370338 | 0.246405 | 0.404661 | 0.411377 |
| CINGHipp | -1.9006 | 0.096505 | 0.166594 | 0.105157 | 0.085073 | 0.060728 | 0.065631 | 0.086895 | 0.068002 | 0.014313 | 0.049874 | 0.077276 | 0.312649 | 0.116428 | 0.92232 | 0.173318 | 0.156618 | 0.127062 | 0.256132 | 0.2293 | -0.01222 |
| FMaj | -5.16363 | 0.204971 | 0.122581 | 0.203382 | 0.152919 | 0.133269 | 0.126915 | 0.215377 | 0.184593 | 0.011221 | 0.103675 | 0.208996 | 0.124764 | 0.093792 | 0.173318 | 0.786461 | 0.311678 | 0.349728 | 0.329825 | 0.203632 | 0.284242 |
| FMin | -12.0934 | 0.419928 | 0.372896 | 0.46767 | 0.3997 | 0.327474 | 0.365904 | 0.493354 | 0.462903 | 0.015212 | 0.204707 | 0.560993 | 0.280685 | 0.431222 | 0.156618 | 0.311678 | 0.860789 | 0.615351 | 0.511213 | 0.47517 | 0.48335 |
| IFOF | -9.14884 | 0.35834 | 0.311892 | 0.380358 | 0.316838 | 0.258725 | 0.281843 | 0.406352 | 0.378202 | 0.033963 | 0.190048 | 0.570611 | 0.317736 | 0.370338 | 0.127062 | 0.349728 | 0.615351 | 0.768242 | 0.595994 | 0.522629 | 0.588434 |
| ILF | -7.82955 | 0.293436 | 0.233897 | 0.310533 | 0.246517 | 0.219337 | 0.238493 | 0.320487 | 0.317319 | 0.024775 | 0.12005 | 0.36261 | 0.305891 | 0.246405 | 0.256132 | 0.329825 | 0.511213 | 0.595994 | 0.792123 | 0.505552 | 0.377276 |
| SLF | -5.91601 | 0.227084 | 0.240043 | 0.259658 | 0.22572 | 0.205932 | 0.23145 | 0.282275 | 0.252321 | 0.023844 | 0.149826 | 0.424691 | 0.372024 | 0.404661 | 0.2293 | 0.203632 | 0.47517 | 0.522629 | 0.505552 | 0.703961 | 0.388659 |
| UNC | -6.31008 | 0.259104 | 0.204627 | 0.308776 | 0.24521 | 0.192378 | 0.208202 | 0.282191 | 0.295929 | -0.02542 | 0.1042 | 0.460745 | 0.195309 | 0.411377 | -0.01222 | 0.284242 | 0.48335 | 0.588434 | 0.377276 | 0.388659 | 0.941481 |

**Supplementary Table 2:** Unstandardized parameter estimates and SE for the model shown in Figure 6. = ∼ denotes factor loadings, ~ denote structural paths.

| Parameter | estimate | se | Parameter | estimate | se |
| --- | --- | --- | --- | --- | --- |
| gf=~cattell1 | 0.50133 | 0.03070 | CRTcons~ATR | 0.10967 | 0.05019 |
| gf=~cattell2 | 0.46154 | 0.02903 | CRTcons~CST | 0.00408 | 0.04258 |
| gf=~cattell3 | 0.50836 | 0.02939 | CRTcons~CING | -0.05019 | 0.04596 |
| gf=~cattell4 | 0.45607 | 0.02765 | CRTcons~CINGHipp | -0.01278 | 0.04102 |
| gf~SRTspeed | -0.05160 | 0.08572 | CRTcons~FMaj | -0.00994 | 0.03940 |
| gf~SRTcons | 0.21473 | 0.07838 | CRTcons~FMin | 0.41584 | 0.06121 |
| gf~CRTspeed | 0.58450 | 0.12388 | CRTcons~IFOF | 0.06772 | 0.09115 |
| gf~CRTcons | 0.45310 | 0.10639 | CRTcons~ILF | 0.08362 | 0.06607 |
| gf~AVspeed | -0.40707 | 0.08521 | CRTcons~SLF | -0.08379 | 0.06589 |
| gf~AVcons | 0.32510 | 0.08505 | CRTcons~UNC | 0.02082 | 0.05591 |
| SRTspeed~ATR | -0.01532 | 0.06102 | AVspeed~ATR | -0.10657 | 0.07467 |
| SRTspeed~CST | 0.13216 | 0.04895 | AVspeed~CST | 0.05214 | 0.05024 |
| SRTspeed~CING | -0.05454 | 0.04841 | AVspeed~CING | 0.11468 | 0.05138 |
| SRTspeed~CINGHipp | -0.04206 | 0.04479 | AVspeed~CINGHipp | -0.01839 | 0.04624 |
| SRTspeed~FMaj | 0.01136 | 0.04509 | AVspeed~FMaj | 0.00912 | 0.04977 |
| SRTspeed~FMin | 0.33476 | 0.06922 | AVspeed~FMin | -0.03167 | 0.07041 |
| SRTspeed~IFOF | 0.03230 | 0.10333 | AVspeed~IFOF | 0.20781 | 0.10469 |
| SRTspeed~ILF | -0.00097 | 0.06929 | AVspeed~ILF | -0.03193 | 0.07308 |
| SRTspeed~SLF | 0.01050 | 0.06227 | AVspeed~SLF | -0.03892 | 0.06643 |
| SRTspeed~UNC | -0.00066 | 0.05990 | AVspeed~UNC | -0.13510 | 0.05845 |
| SRTcons~ATR | 0.06517 | 0.05725 | AVcons~ATR | -0.01377 | 0.06324 |
| SRTcons~CST | 0.10191 | 0.04260 | AVcons~CST | 0.05797 | 0.04756 |
| SRTcons~CING | -0.06104 | 0.04939 | AVcons~CING | 0.05048 | 0.05126 |
| SRTcons~CINGHipp | -0.03547 | 0.04159 | AVcons~CINGHipp | -0.00933 | 0.04669 |
| SRTcons~FMaj | -0.01233 | 0.04430 | AVcons~FMaj | 0.04584 | 0.04870 |
| SRTcons~FMin | 0.36707 | 0.06678 | AVcons~FMin | 0.13174 | 0.06671 |
| SRTcons~IFOF | -0.01815 | 0.09662 | AVcons~IFOF | 0.24800 | 0.09819 |
| SRTcons~ILF | 0.01698 | 0.06497 | AVcons~ILF | -0.14182 | 0.07040 |
| SRTcons~SLF | 0.02887 | 0.06062 | AVcons~SLF | 0.03291 | 0.07023 |
| SRTcons~UNC | -0.00483 | 0.05895 | AVcons~UNC | -0.12571 | 0.05655 |
| CRTspeed~ATR | 0.06524 | 0.04813 |  |  |  |
| CRTspeed~CST | 0.10411 | 0.04112 |  |  |  |
| CRTspeed~CING | -0.14056 | 0.04629 |  |  |  |
| CRTspeed~CINGHipp | -0.02453 | 0.04053 |  |  |  |
| CRTspeed~FMaj | 0.02173 | 0.03944 |  |  |  |
| CRTspeed~FMin | 0.47897 | 0.06044 |  |  |  |
| CRTspeed~IFOF | 0.17490 | 0.08824 |  |  |  |
| CRTspeed~ILF | -0.04620 | 0.06300 |  |  |  |
| CRTspeed~SLF | -0.02702 | 0.06140 |  |  |  |
| CRTspeed~UNC | -0.04108 | 0.05723 |  |  |  |

## Appendix A Estimating WM ROIs

Diffusion-weighted images (DWI) were acquired at the MRC Cognition and Brain Sciences Unit, using a 3T Siemens TIM Trio MRI scanner, and a 32-channel head coil. A twice-refocused spin-echo sequence was used to minimise eddy-currents, with 30 uniformly spaced gradient directions for each of two b-values (1000 and 2000 s/mm2), and three non-diffusion weighted images (b-value = 0). Other imaging parameters were: TR=9100ms, TE=104ms, 2x2x2mm3 resolution, FoV 192x192mm2, 66 axial slices and GRAPPA acceleration factor of 2. A structural MPRAGE was also acquired for each participant (see Shafto et al. (2014) for sequence details). Traditionally, DWI data is motion corrected at the postacquisition level by using image registration techniques to co-register each diffusion-weighted image to the first acquired b=0 image. However, as discussed in Ben-Amitay et al. (2012), when high b-values are used, this technique will fail to correct for distortions and motion, and may introduce other artefacts. For this reason, we did not apply registration-based motion correction to the DWI data in this study. Detection and exclusion of outliers was performed at the analysis level to avoid including datasets affected by other artefacts and distortions. All pre-processing and modelling of MRI data was performed using a combination of functions from FSL version 5.0.8 (Jenkinson, Beckmann, Behrens, Woolrich, & Smith, 2012), SPM12 (http://www.fil.ion.ucl.ac.uk/spm/software/spm12/, and custom scripts written in C and Matlab, integrated in the Automatic Analysis (aa)package (Cusack et al., 2014). After removal of non-brain tissue, a non-linear diffusion tensor model was applied to the DWI data. Non-linear fitting of the diffusion tensor provides better noise modelling than standard linear model fitting, resulting in more accurate and un-biased estimates of the diffusion tensor and its different metrics (Jones & Basser, 2004). The diffusion tensors eigensystem was used to compute the fractional anisotropy (FA) at each voxel. The FA maps were then spatially normalised into a standard stereotactic space as follows: firstly, the average of the three b=0 images for each subject was coregistered to their MPRAGE; secondly, the MPRAGE images were coregistered across participants to a sample-specific template using DARTEL (Ashburner, 2007), and the transformations so derived were used to warp each participant's images into standard MNI space, including their FA maps. The resulting FA images in MNI space were smoothed with a 1mm FWHM Gaussian kernel to reduce residual interpolation errors.

---

**Supplementary Figure 1.**
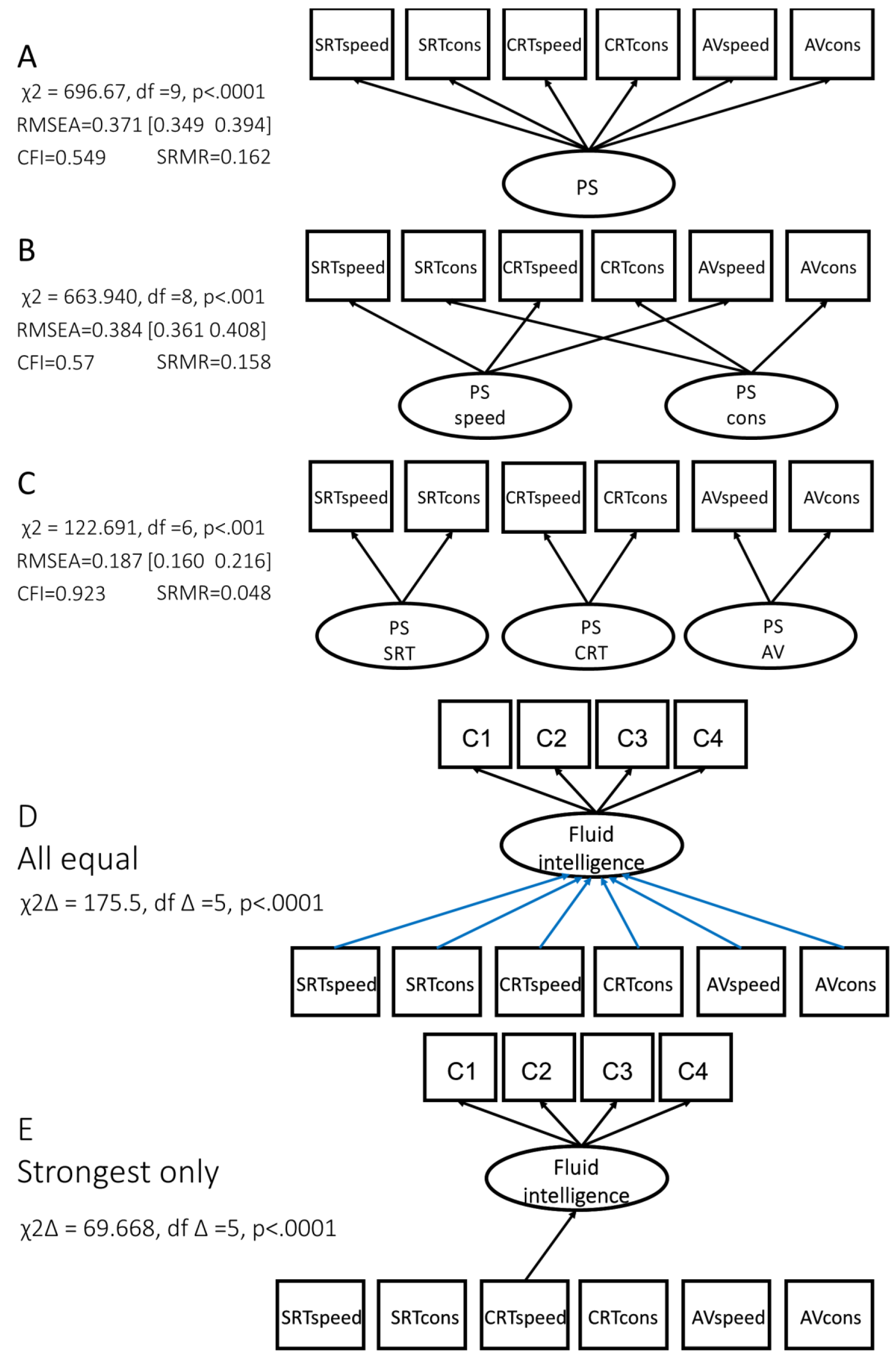
Competing models for processing speed (A-E). A: unidimensional model of PS. B: Separate latent variables for speed and consistency. C: A latent variable for each task. D: An equality constrained model, where the effect of each PS variable is constrained to be equal. E: A model where only the strongest effect is estimated freely, all others constrained to be zero. All models fit worse than the freely estimated model reported in the text. For models D and E, residual covariance between PS variables is allowed but not shown for clarity.

**Supplementary Figure 2.**
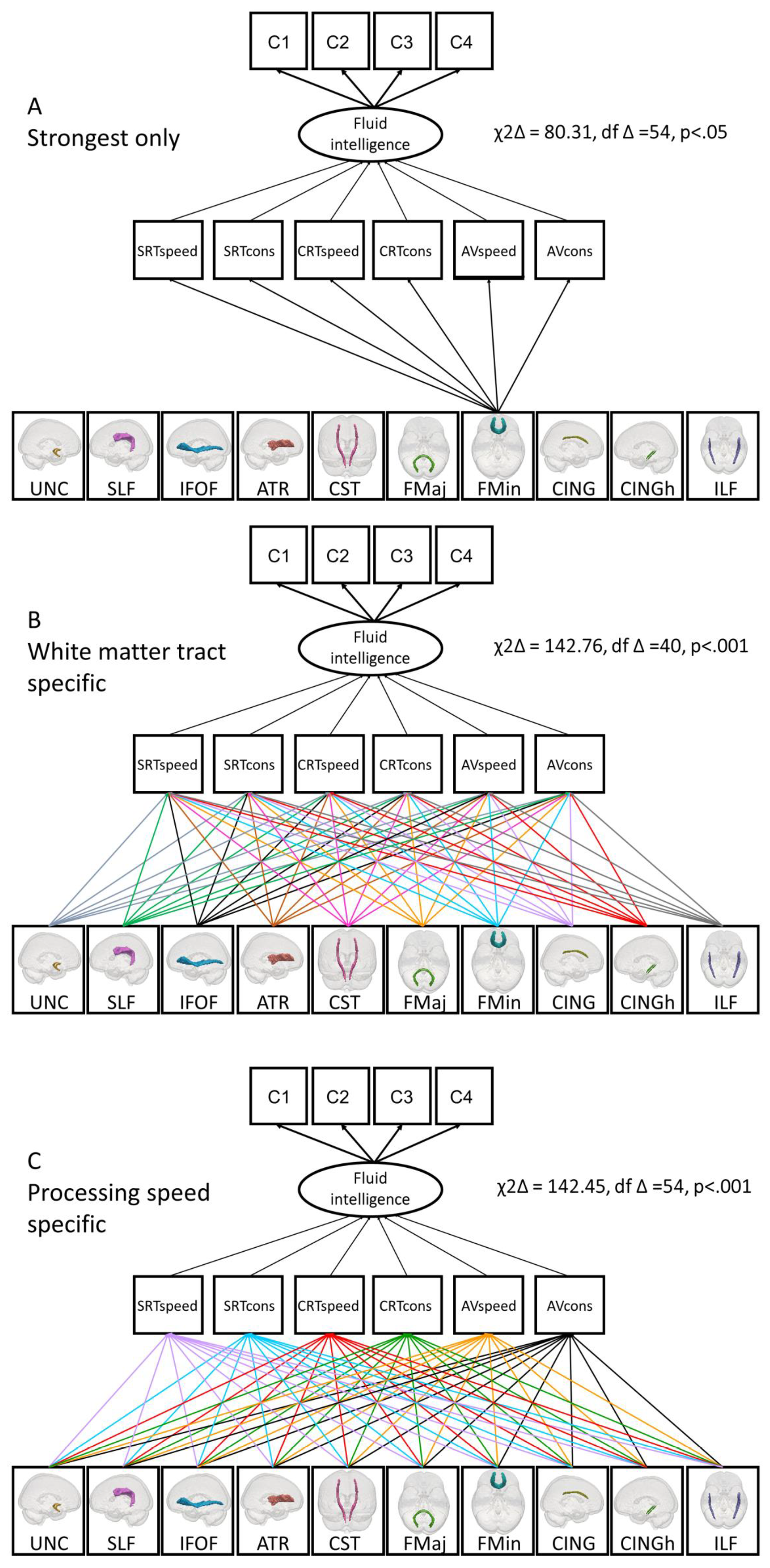
Competing models for the relationship between white matter integrity and processing speed. A: ‘Strongest only’ model, where the strongest standardized path (Forceps minor to PS) is estimated freely, but all others constrained to be zero. B White matter tract specific model: the effect of each WM tract is constrained to be equal across PS, but can vary between tracts. C: Processing speed specific model: The effect of white matter is allowed to vary across processing speed domains, but presumed to be equal for each tract. All models fit worse than the freely estimated model reported in the text. Note: Residual covariance between PS variables and WM tracts are allowed but not shown for clarity.

## Footnotes

1 Note that this notion of a ‘watershed’ should not be confused with the term ‘watershed’ as used in reference to differential cerebral blood perfusion and arterial beds in aging, e.g. Raz, (2005); Suter et al., (2002).

